# The Effect of Choline Alphoscerate on Non spatial memory and Neurogenesis in a Rat Model of Dual Stress

**DOI:** 10.1101/2020.06.16.154310

**Authors:** Hyo Jeong Yu, Min Jung Kim, Jung Mee Park, So Young Park, Shi Nae Park, Dong Won Yang

## Abstract

Choline alphoscerate (α-GPC) is a choline-based compound and acetylcholine precursor commonly found in the brain; it has been known to be effective in treating neuronal injury and increasing the levels of acetylcholine (Ach) and brain-derived neurotrophic factor (BDNF) which in turn enhances memory and cognitive function. This study was designed to establish rat models of dual stress using noise and restraint in order to investigate the effect of α-GPC on cognitive function and neurogenesis after dual stress. The rats were randomly divided into four groups as follows: a control group (CG), a control with α-GPC group (CDG), a noise-restraint stress group (NRSG), and a noise-restraint stress with α-GPC group (NRSDG). Two experimental groups were exposed to the double stress stimuli of noise and restraint, which involved 110dB sound pressure level (SPL) white band noise and restraint at the same time for 3 hours/day for 7 days. While the CG and NRSG received saline, the CDG and NRSDG received α-GPC (400mg/kg) orally after stress exposure. The α-GPC–treated group showed increased memory function compared to the dual stress group in the novel object recognition test. In analysis of the hippocampus, the α-GPC–treated group showed greater Choline acetyltransferase (ChAT) and BDNF expression compared to the dual stress group. The α-GPC–treated group showed significantly increased neuroblast expression compared to the dual stress group, which suggests that α-GPC enhances BDNF expression and protects the activity of the immature cells at the dentate gyrus. Our results suggest that α-GPC treatment can protect cognitive function and neurogenesis in a dual stress model.

## 1. Introduction

Certain physical or psychological stressors disrupt the homeostasis of animals (1, 2). Noise stress can cause damage to cochlear hair cells, which induces hearing loss, and it also impairs non-auditory systems that caused cognitive dysfunction, sleep disturbance as well as physical changes such as neurotransmitters and immune system. Restraint stress in rodents impairs their feeding behavior and emotions, and it is also known to be the most extreme stressor to occur behavioral, neurochemical and immunological changes in response to various types of stress (3, 4).

The brain plays the main role in recognizing the intensity of sound exposure, and is involved in interpreting and responding to potential stressors (5). In the adult brain, the subgranular zone (SGZ) of the hippocampus plays an important role in memory and learning as a major site of neurogenesis (6). The hippocampus is vulnerable to neurotoxic conditions and factors such as stress and depression, which have harmful effects on neurogenesis (7). Studies have been shown chronic restraint stress caused loss of hippocampal cell and reduction of hippocampus neurogenesis (8).

Brain-derived neurotrophic factor (BDNF) is a member of the family of nerve growth factors, which are otherwise known as neurotrophins. BDNF is involved in various functions, ranging from food intake to behavior, spatial and non-spatial memory associated with central nervous system structures such as the hippocampus, cerebral cortex, and hypothalamus (9, 10). BDNF in the hippocampus is known to induce neuronal development by promoting the normal development, survival and plasticity of neurons, and differentiation of SGZ progenitor cells (11). However, experimental studies showed that stress can decrease BDNF expression in the hippocampus (12).

Choline alphoscerate (α-GPC) is a choline-based compound commonly found in the brain and is an important intermediate in the synthesis of both acetylcholine (Ach) and cell membrane phospholipids as a cholinergic precursor (13). Increasing the Ach levels in the hippocampus is known to enhance cholinergic neurotransmission and to increase learning and memory by enhancing BDNF expression (7, 14–16). Studies have been shown that α-GPC is effective in enhancing cognition in animal models of Alzheimer’s disease or dementia (17). However, the effect of α-GPC on the severely stressed brain has, to the best of our knowledge, never been studied before. The purpose of this study was to induce dual stresses in rats using noise and restraint, and to investigate the effect of α-GPC on memory function as well as neurogenesis and neuronal protection in the hippocampus after severe stress.

## 2. Materials and methods

### 2.1. Animals and Grouping

Eight-week-old male Wistar rats (240–320g) were purchased from Orient Bio (Sungnam, Korea). The rats were randomly divided into 4 age-matched groups as follows: a control group (CG); a control with α-GPC drug administered group (CDG); a noise and restraint stress group (NRSG); a noise and restraint stress with α-GPC drug administered group (NRSDG).

All procedures for animal research were performed in accordance with the Laboratory Animals Welfare Act, the Guide for the Care and Use of Laboratory Animals, and the Guidelines and Policies for Rodent Experiments provided by the IACUC (Institutional Animal Care and Use Committee) in the College of Medicine, The Catholic University of Korea. All animals were kept in a pathogen-free environment on a 12-h light/dark cycle and had access to gamma-ray sterilized food (TD 2018S, Harlan Laboratories, Inc., Indiana, USA) and autoclaved water ad libitum.

### 2.2. Induction of stress and drug administration

Two of the experimental groups were exposed to the dual stress stimuli of noise and restraint at the same time. Rats in a pie-shaped wire cage were exposed to 110 dB sound pressure level (SPL) white band noise, and restraint stress was administered by putting the rats in cylindrical plastic films, DecapiCones, with rubber bands fixed at the tails for 3 hours/day for 7 days. While the CG and the NRSG received saline, the other 2 groups received α-GPC (400mg/kg) orally immediately after the dual stress exposure (**Fig 1**).

**Figure 1.**
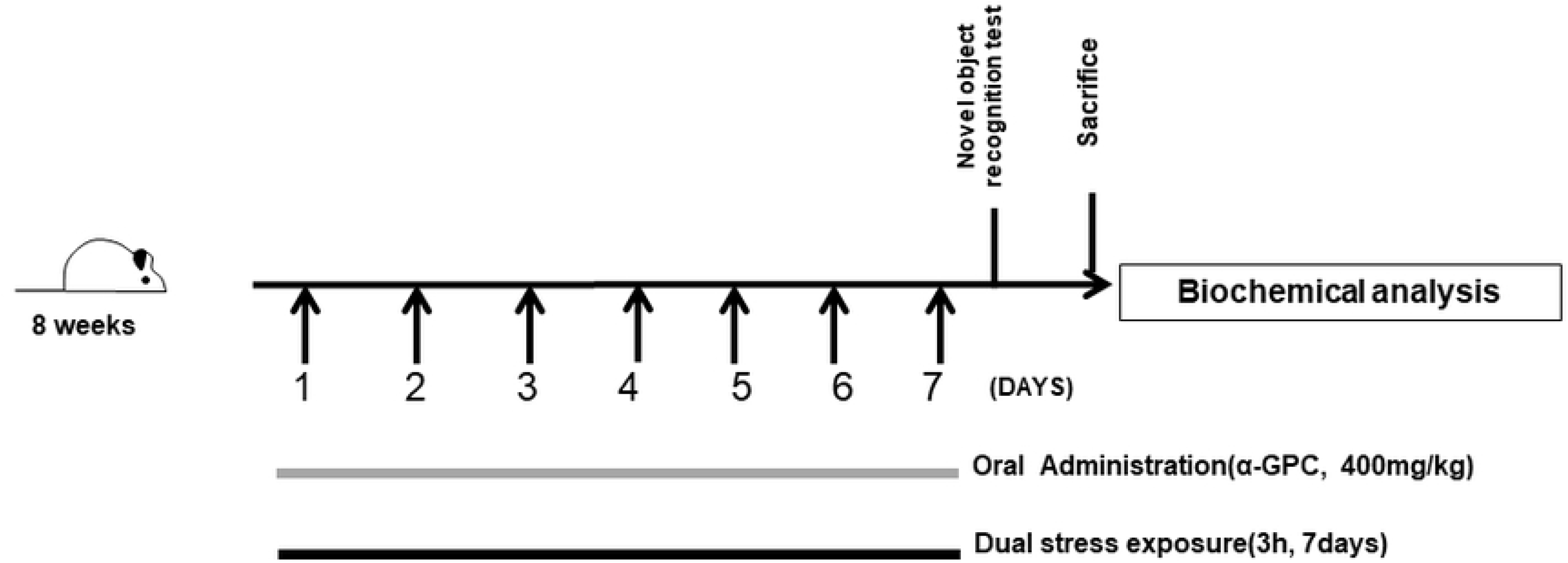
Experimental design.

### 2.3. Auditory brainstem response and noise exposure

All hearing tests were performed with the rats under anesthesia, using a mixture of Rompune (0.4 ml/kg) and Zoletil (0.6 ml/kg). The auditory brainstem response was recorded using an Intelligent Hearing System (IHS) Smart EP fitted with high-frequency transducers (HFT9911-20-0035) and running IHS high-frequency software version 2.33 (IHS, Miami, FL). The details of the hearing test were described in our previous study (18). ABR thresholds and DPOAE levels were compared among the 4 groups in this study.

### 2.4. Morphological measures of the organ of Corti by light microscopy

The dissected cochleae were perfused through the round and oval windows with 0.1 M phosphate buffer containing 2 % paraformaldehyde and 2 % glutaraldehyde, and incubated in the same fixatives overnight at 4 °C. They were then rinsed with 0.1 M PBS, perfused in 1 % osmium tetroxide, and incubated overnight, followed by immersion in 0.5 M EDTA, which was changed every day for 2 weeks. The decalcified cochleae were dehydrated in 50, 70, 90, 95, and 100 % ethanol and 13.5 M acetone, and embedded in Araldite 502 resin (Electron Microscopy Sciences, Fort Washington, PA, USA). The cochleae were sliced into 5-um sections, stained with toluidine blue, and mounted in Permount on microscope slides. The slides were scanned using a digital microscopy scanner (Pannoramic MIDI, 3DHISTECH Ltd., Budapest, Hungary) with a 20× microscope objective. The computer software Pannoramic Viewer 1.15.2 (3DHISTECH Ltd., Budapest, Hungary) was used for image viewing. Three different regions of the cochlea (apex, middle, and base) from the mid-modiolar sections with a clearly visible whole organ of Corti (OC) were chosen for image magnification (X35 objective). A modified rank-order grading method was used to rate the status of the OC. Numbers were assigned to indicate the following conditions: complete degeneration 1); a low cuboidal cell layer without recognizable supporting cells 2); partial collapse of the OC, but with subtypes of supporting cells still recognizable 3); maintenance of normal cytoarchitecture of the OC with all supporting cells intact, but loss of hair cells 4); and normal cytoarchitecture of the OC with intact hair cells 5). The averaged regional scores for the OC in the base, middle, and apical turns were compared between the 4 groups.

### 2.5. Cognitive behavioral test

Non-spatial memory was assessed with the novel object recognition (NOR) test, based on the experimental protocol described earlier with a slight modification (2). Simply, a 60 × 60 cm, grey-colored, polyvinyl chloride plastic box was used. The whole test period consisted of 4 sessions: handling, habituation, adaptation, and testing. After each rat was handled for 20 min, it was placed in the apparatus to freely explore the environment (15 min/day, 3 days). On the next day, each rat was placed in the apparatus facing the wall opposite to the segment in which 2 identical sample objects were placed. The rat was immediately put back into its home cage after freely exploring these objects for 10 min. After 1h, the test was performed. One of the sample objects was replaced by a novel object, and the rat was placed in the same way. Exploration and recognition of the objects were defined as the nose being in contact with an object or directed to the object within a defined distance (<2 cm). Sitting or leaning on the object was not included in the definition of exploration or recognition. After assessment of each animal, the apparatus was properly cleaned with 70 % ethanol to prevent olfactory cues or stimuli. Behaviors of animals were recorded with a video camera placed at the top of the box, and we used the XNote Stopwatch program to record the amount of time the rats explored.

### 2.6. Staining of the hippocampus

Rats were transcardially perfused with saline solution containing 0.5 % sodium nitrate and heparin (10 U/ml), and then it fixed with 4 % paraformaldehyde (PFA) in phosphate buffer saline (PBS). The hippocampus was dissected, post-fixed overnight in buffered 4 % paraformaldehyde at 4 °C, PBS washed, and stored in a 30 % sucrose solution at 4 °C until it sank. Free-floating sectioning was performed at −20 °C on a Cryostat Microtome (Leica, Germany) to generate 30 μm-thick coronal sections. Sections were stored at −20 °C in cryoprotect solution until use. For immunofluorescence staining, brain sections were then washed with PBS and incubated for 24 h at 4 °C with anti-BDNF (Abcam, USA, 1:400), anti-doublecortin (DCX) (Doublecortin, identify immature neurons; Cell Signaling Technology, USA) and NeuN (Fox-3, Millipore, USA) antibody, diluted 1:100 with 0.5 % BSA in PBS.

The sections were then washed in PBS and subsequently incubated with Alexa 488 goat antirabbit (1:200, Thermo Fisher Scientific Inc., USA) or Alexa 555 goat anti-mouse (1:200, Thermo Fisher Scientific Inc., USA) antibody for 2 h in the dark. Sections were washed thoroughly and DAPI mounted with fluorescent mounting medium (Vector Laboratories, Burlingame, CA, USA). Nuclei were counterstained with blue-fluorescent DAPI. Stained sections were examined by confocal laser scanning microscopy (LSM 800 Meta, Carl Zeiss, Jena, Germany). An experimenter blinded to group counted the number of DCX-positive cells and the intensity of BDNF in the SGZ of the dentate gyrus (DG) from one hemisphere at each 200x, 400x magnification. A series of 3 z-stack images were acquired per hippocampal field at 1.5 μm intervals under a magnification of X400 for BDNF and A series of 6 z-stack images were acquired per hippocampal field at 1.8 μm intervals under a magnification of X200 for DCX. A single digital image was reconstructed from the z-stacks using ZEN 2012 software (Carl Zeiss, Jena, Germany).

For immunohistochemistry staining, brain sections were washed with PBS and incubated with 3% H2O2 for 10 m. sections were washed with PBS for 2 times and incubated with 1.5 % BSA in PBST for 1 h. After blocking step, sections were incubated with anti-Interlukin-1β (IL-1β, Abcam, 1:500) and anti-Choline acetyltransferase (ChAT, Milipore, 1:500) with 0.5 % BSA in PBS for 24 h at room temperature. After washing with 0.5 % BSA for 2 times, sections were incubated for 2 h at room temperature with biotinyl rabbit antibody (Vector Laboratories, 1:1000) in 0.5 % BSA. After washing with 0.5 % BSA for 2 times, brain sections were incubated with Avidin–Biotin Complex (ABC) reagent (Vector Laboratories) in PBS for 2 h. After washing with PBS for 2 times, Sections were developed using diaminobenzadine (DAB) peroxidase substrate kit (Vector Laboratories) and mounted on slides, air-dried, and cover slipped for microscopic observation. The number of positive neurons in the IL-1β and ChAT was counted in the SGZ of the DG from one hemisphere at 400x magnification using a microscope rectangle grid.

### 2.7. Assessment of the plasma corticosterone level

The plasma level of the stress hormone corticosterone was also evaluated. After exposure to the dual stress for 7 days, rats were deeply anesthetized and blood was quickly collected via the retro-orbital plexus. Blood was collected in a heparinized Eppendorf tube and immediately centrifuged at 13,000 rpm for 15 mins at 4 °C to separate the plasma, which was stored at −80 °C until use. Corticosterone concentration was measured by a competitive enzyme-linked immunoassay (Corticosterone competitive ELISA kit, Abcam, USA) according to the manufacturer’s protocol. The optical density was measured at 450 nm using an ELISA reader.

### 2.8. Histological assessment of hippocampus

Rats were transcardially perfused with saline solution containing 0.5 % sodium nitrate and heparin (10 U/ml), and then fixed with 4 % PFA in PBS. Each brain was dissected, post-fixed overnight in buffered 4 % PFA at 4 °C, PBS washed, and stored in a 30 % sucrose solution at 4 °C until it sank. Coronal brain sections including hippocampus were sliced to a thickness of 10 μm, and were stained with hematoxylin and eosin (H&E). Neuronal cell numbers were counted using NIH ImageJ software.

### 2.9. Data analysis

Statistical significance was assessed by comparing mean ± standard error of the mean (SEM) values using the Student’s t-test for single comparisons or analysis of variance (ANOVA) followed by Tukey’s HSD, LSD post-hoc test for multiple comparisons.

## 3. Results

### 3.1. Functional and histological changes of auditory system after dual stress

#### 3.1.1. Hearing tests

The ABR threshold and the DPOAE levels were measured to confirm that they were properly exposed to noise stress. After dual stress exposure, the NRSG and NRSDG showed hearing loss, with mean ABR thresholds of 56.25 ± 4.41, 61.25 ± 5.77, 71.25 ± 11.66, and 81.25 ± 3.33 dB sound pressure levels (SPL) and of 63.75 ± 2.88, 63.75 ± 5.00, 80 ± 10.40, and 83.75 ± 6.00 dB SPL for click, 8kHz, 16kHz, and 32kHz respectively, which were significantly higher hearing threshold levels in the stress group than in the control group (p < 0.001 for 8, 16, click kHz; p < 0.05 for 32 kHz) (**Fig 2A**). The DPOAE levels were also significantly different between the experimental group and the control group at the 6 to 14 kHz geometric mean (GM) frequencies (kHz) (**Fig 2B**). At the higher GM frequencies, DPOAE responses were small in all groups and there was significance at only 26 kHz GM.

**Figure 2.**
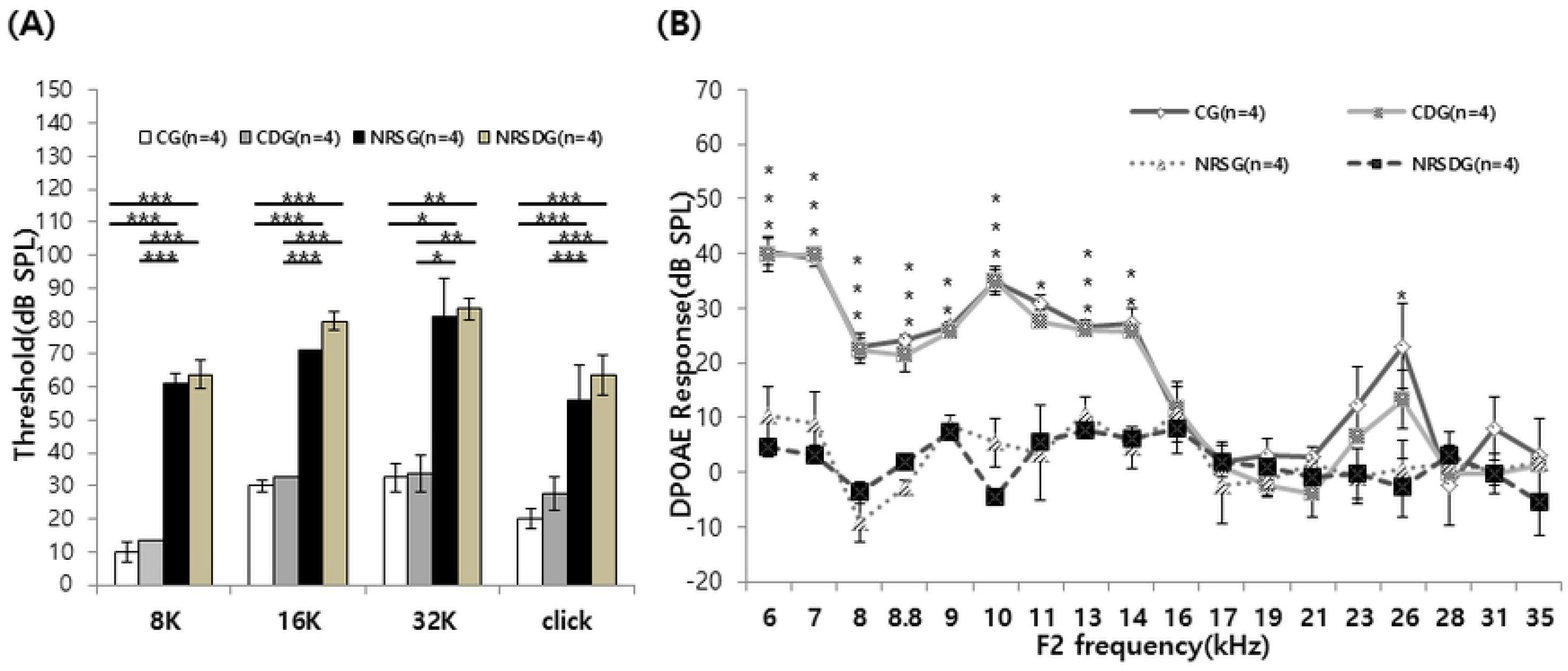
Hearing test results: auditory brainstem response thresholds for click and 8/16/32 kHz tone bursts and distortion product optoacoustic emission (DPOAE) at 6 to 32kHz in rats. ABR thresholds in all groups after the dual stress of noise and restraint exposure. The stress groups (NRSG, NRSDG) showed increased ABR thresholds at all kHz compared to the control groups (A). The stress groups showed significantly decreased DPOAE responses compared to the control groups in outer hair cell function at the 6 to 14kHz low-frequency regions and 26kHz after dual stress (B). (Error bars indicate SEM, ANOVA, Tukey’s HSD post-hoc, * p< 0.05, ** p< 0.01, *** p< 0.001) (N=4).

#### 3.1.2. Organ of Corti (OC) grading

The grades for OC degeneration in the base, middle and apex in the control group were 4.61 ± 0.05, 4.61 ± 0.03, and 4.71 ± 0.04, respectively. Those in the CDG were 4.55 ± 0.04, 4.48 ± 0.04, and 4.53 ± 0.07, respectively. Those in the NRSG were 3.46 ± 0.17, 3.54 ± 0.17, and 3.41± 0.22, respectively. Those in the NRSDG were 3.23 ± 0.13, 3.55 ± 0.10, and 3.55 ± 0.17, respectively (**Figs 3A and 3B**). The grading scores for the organ of Corti for NRSG and NRSDG were significantly decreased compared to the CG at all cochlear turns (p < 0.01).

**Figure 3.**
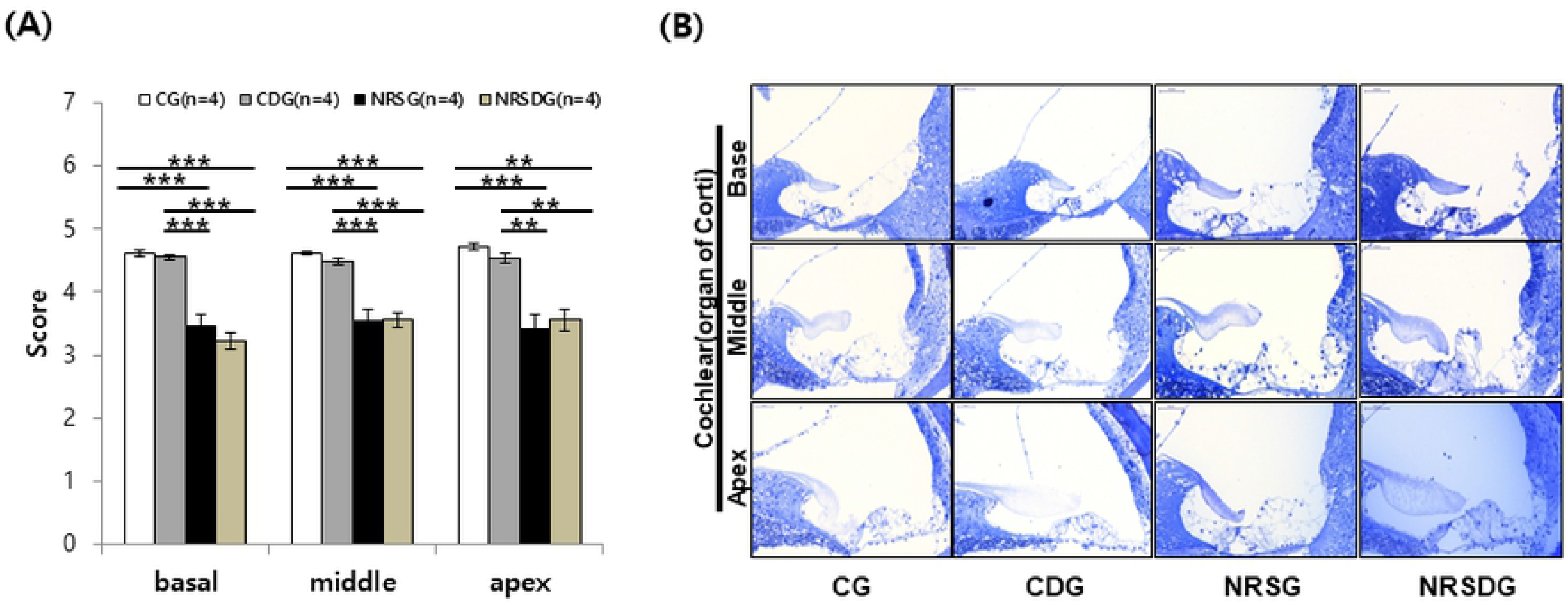
Light microscopic morphology of the organ of Corti. Representative photomicrographs of the organ of Corti from the basal, middle, and apical turns. The dual stress groups (NRSG, NRSDG) showed severely damaged and collapsed morphology (magnification X350) (A). The stress groups demonstrated significantly decreased grading scores of the organ of Corti compared to the control groups at the basal, middle, and apical turns. (Error bars indicate SEM, ANOVA, Tukey’s HSD post-hoc, ** p< 0.01, *** p< 0.001) (N=4).

### 3.2. Physiological changes

#### 3.2.1. Body weight

To investigate the effect of dual stress on feeding behavior, we measured the body weight of each rat on the next day after 7 days of dual stress. The CG, CDG, NRSG, and NRSDG mean body weights were 270 ± 2.66 g, 272.5 ± 6.84 g, 230 ± 4.61 g, and 240.3 ± 5.74 g, respectively. The dual-stressed groups showed a greater than 10 % reduction in body weight compared to the CG and CDG (p < 0.05) (**Fig 4A**).

#### 3.2.2. Plasma corticosterone

Corticosterone level has been used as a representative stress marker. The plasma corticosterone concentration was measured by ELISA in this study. The levels of corticosterone in the CG, CDG, NRSG, and NRSDG groups were 540 ± 66.84 ng/ml, 526 ± 83.32 ng/ml, 980 ± 141.73 ng/ml, and 915 ± 57.47 ng/ml, respectively. The plasma corticosterone level in the dual stressed NRSG group was significantly higher than those of the CG and CDG (p < 0.05). However, there were no significant differences in corticosterone levels among the NRSDG and the CG and CDG (p < 0.05) (**Fig 4B**).

### 3.3. Dual stress caused neuronal damage in the hippocampus

#### 3.3.1. Dual stress caused neuronal cell loss

H&E staining was performed to confirm the number of neuronal cells in the hippocampus (**Fig 5A**). The number of neuronal cells in the CG, CDG, NRSG, and NRSDG were 1661 ± 21.8, 1712.33 ± 88.3, 1343.66 ± 20.3, and 1712.33 ± 68.693, respectively. The total neuronal cell number in the DG significantly decreased in the NRSG compared to all the other groups (p < 0.05 for NRSG vs CG, NRSDG). However, the NRSDG showed a significant increase in total neuronal cell numbers compared to the CG and the CDG (p< 0.05) (**Fig 5B**).

**Figure 4.**
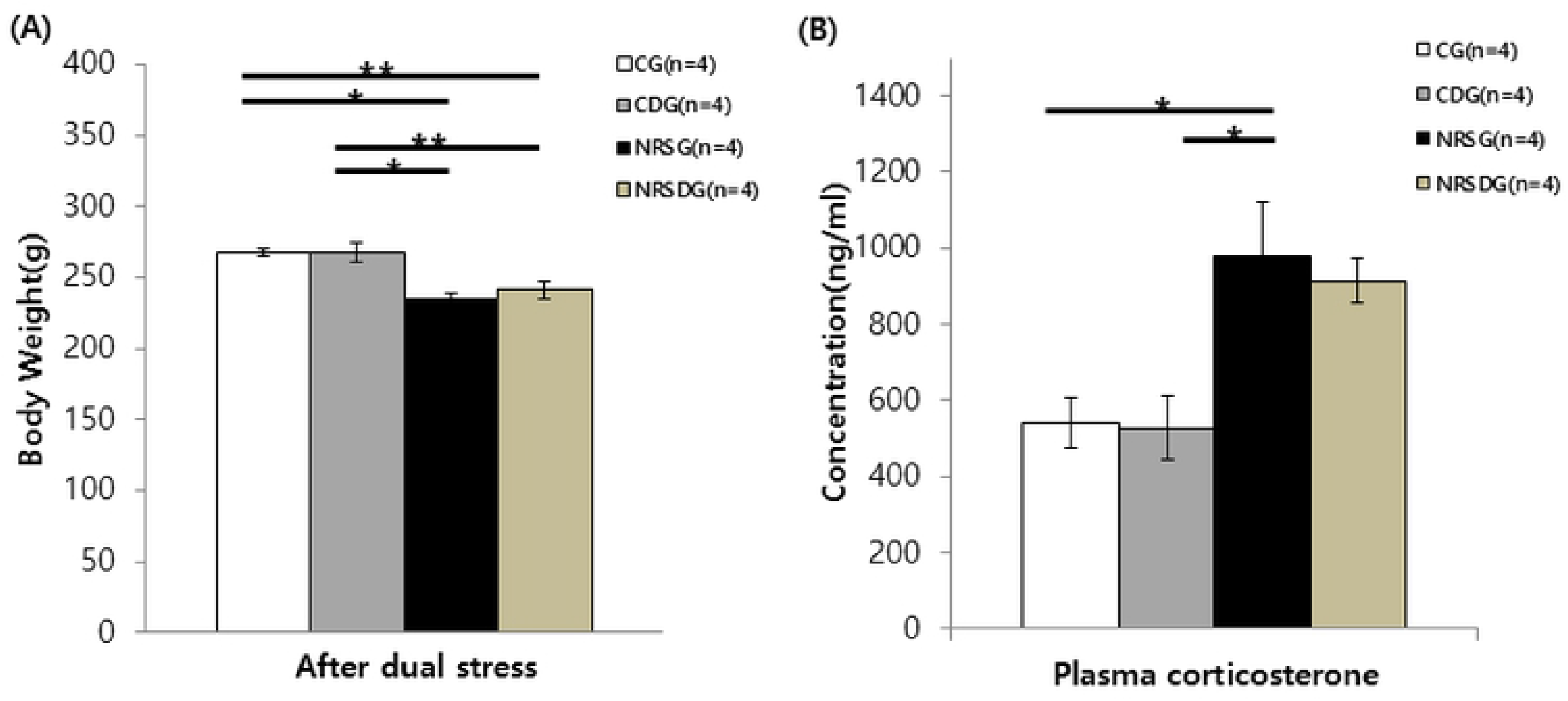
Physiological changes after dual stress. Body weight was measured after dual stress. The dual stress groups (NRSG, NRSDG) showed significantly decreased body weight compared to the control groups, due to stress-induced change in feeding behavior (A). Plasma corticosterone levels were evaluated by ELISA. Significantly increased plasma corticosterone levels were observed in the NRSG compared to the control groups (B). (Error bars indicate SEM, ANOVA, Tukey’s HSD post-hoc, * p< 0.05, ** p< 0.01) (N=4).

**Figure 5.**
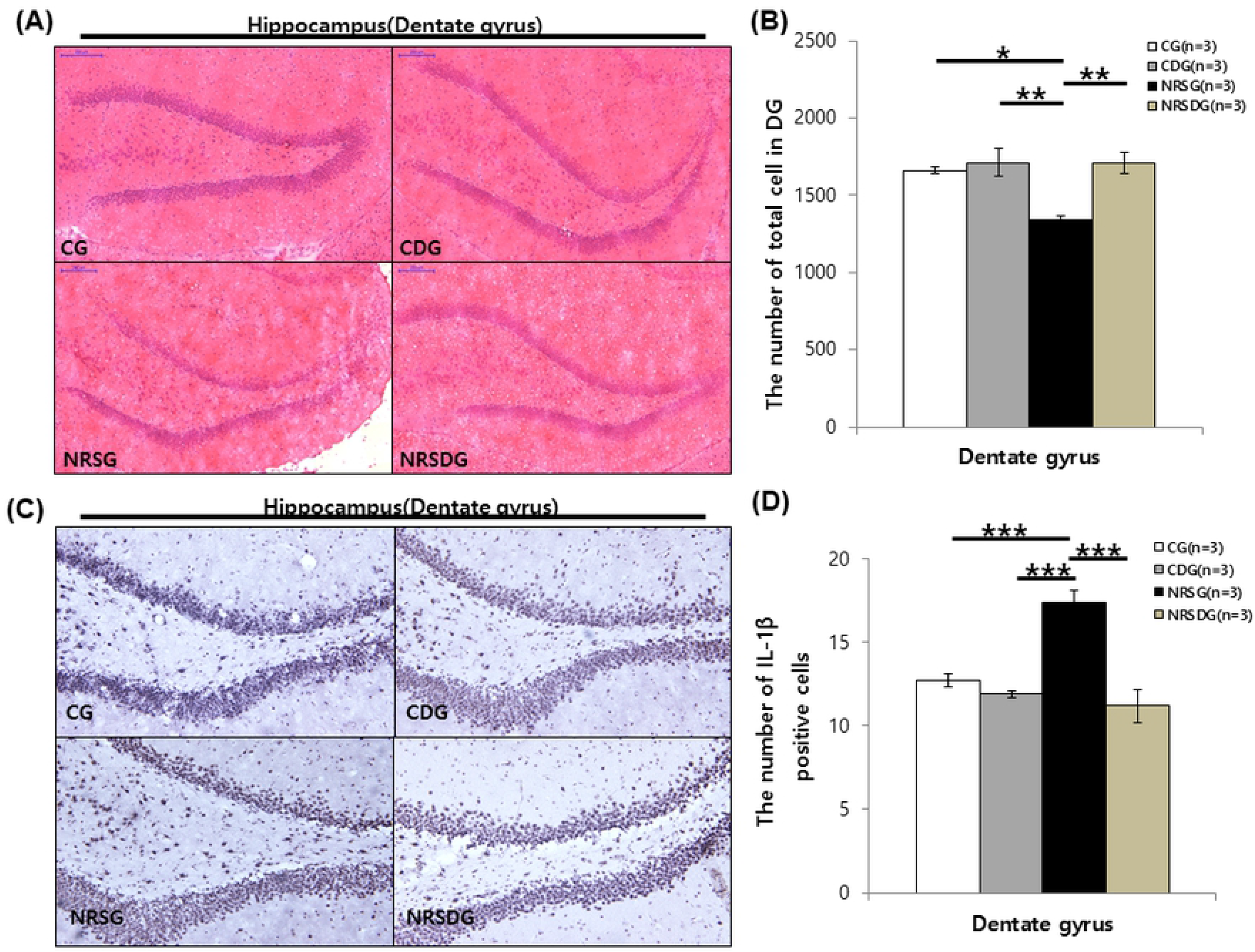
Effect of α-GPC on neuronal loss in hippocampus, histopathological study with hematoxylin and eosin and immunohistochemistry staining. Representative image of dentate gyrus in hippocampus (magnification X70) (A). Alpha-GPC significantly increased the total number of neuronal cells in the NRSDG compared with those in the control groups. The NRSG showed a significantly decreased number of neuronal cells compared to the control groups and the NRSDG (B). Representative image of dentate gyrus in hippocampus (magnification X200) (C). Alpha-GPC significantly decreased the number of positive cells in the NRSDG compared with those in the NRSG. The NRSG showed a significantly increased the number of positive cells compared to all of the groups (D). (Error bars indicate SEM, ANOVA, Tukey’s HSD post-hoc, * p< 0.05, ** p< 0.01, *** p< 0.001) (N=3).

#### 3.3.2. Alpha-GPC affects immune response in the hippocampus

Staining for IL-1β was performed to confirm neuro-immune response in the hippocampus (**Fig 5C**). The number of IL-1β positive cells in the CG, CDG, NRSG, and NRSDG were 12.86 ± 0.13, 12.73 ± 1.63, 18.86 ± 0.85 and 11.66 ± 0.85, respectively. The positive cell numbers in the DG significantly increased in the NRSG compared to all the other groups (p < 0.001) (**Fig 5D)**.

### 3.3. Novel object recognition test

All throughout the adaptation phase, all groups of the rats consumed equal time exploring identical items (left and right). They showed no significant differences in time spent exploring both in the adaptation phase (**Fig 6A**). However, they spent significantly different amounts of time exploring a novel object except for the NRSG group in the test phase (t-test, p< 0.05) (**Fig 6B**). Based on test phase exploration time, the discrimination index (DI) was calculated. The DI of the CG, CDG, NRSG, and NRSDG were 47.35 ± 4.07, 49 ± 6.09, 17.06 ± 7.22, and 38.85 ± 3.68, respectively. The NRSG showed a significantly decreased DI compared to the other groups, and the NRSDG showed an increased DI compared to the NRSG (p< 0.05) (**Fig 6C**).

**Figure 6.**
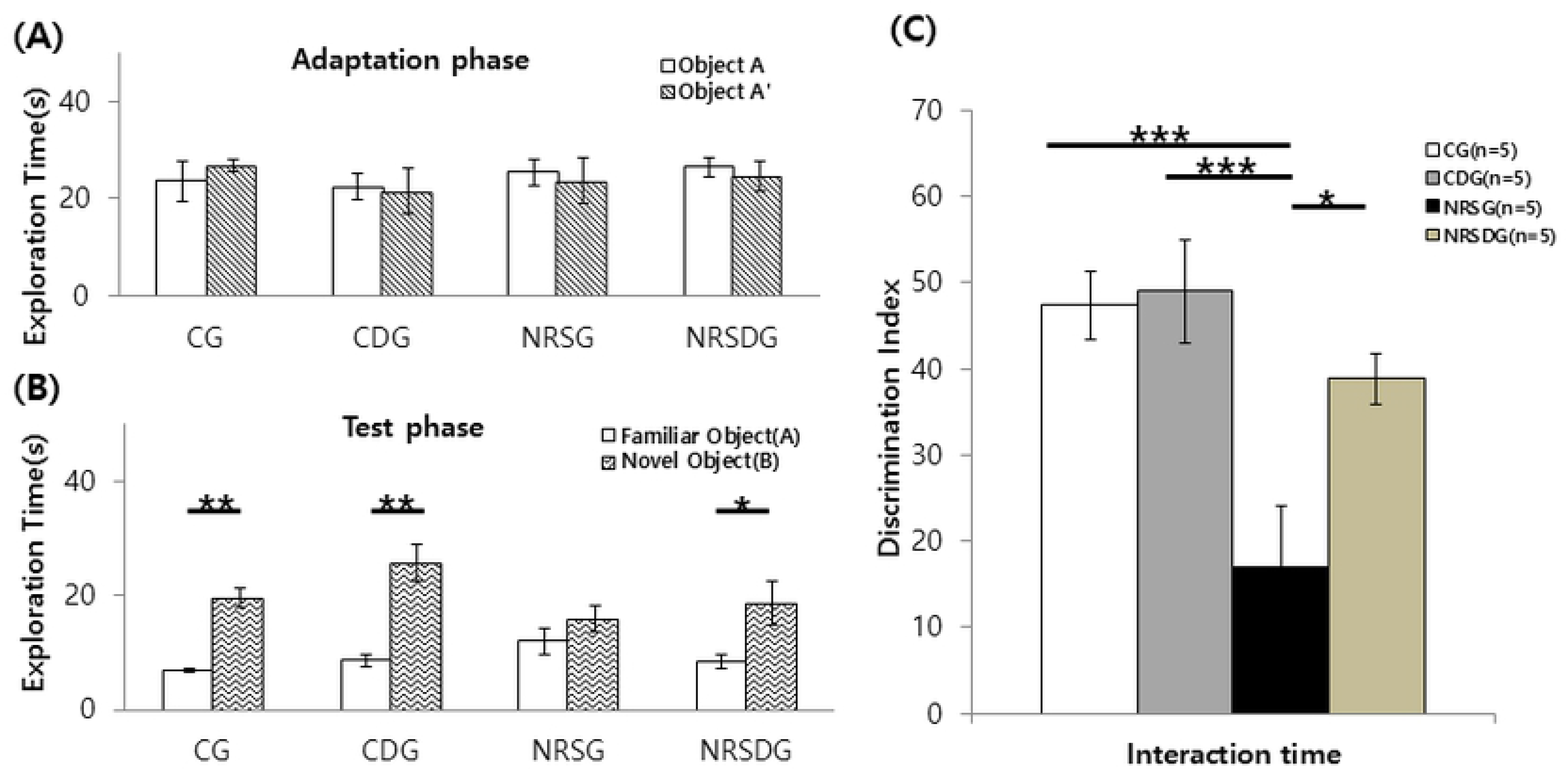
The novel object recognition (NOR) test for cognitive function, measuring the time spent in object exploration during the adaptation and test phases. There were no significant differences among the groups in the adaptation phase, but the rats in the control groups and the NRSDG explored novel objects more frequently than familiar ones in the test phase. The rats in the NRSG did not show a difference in exploration time for novel objects, which indicates decreased cognitive function after dual stress (Student’s t-test) (A&B). The discrimination index (DI) results demonstrated that the rats exposed to dual stress failed to distinguish familiar objects from novel ones. The NRSG showed significantly decreased DI compared to the control groups and the NRSDG (C). (Error bars indicate SEM, Student’s t-test, ANOVA, Tukey’s HSD post-hoc, * p< 0.05, ** p< 0.01, *** p< 0.001) (N=5).

### 3.4. Assessment of ChAT expression for Ach levels in the hippocampus

Immunohistochemistry was performed to evaluate the effect of α-GPC on ChAT expression in the hippocampus (**Fig 7A**). The number of positive cells ChAT expression in CG, CDG, NRSG, and NRSDG were 29.65 ± 0.36, 30.62 ± 0.56, 21.37 ± 0.81, and 29.62 ± 0.43, respectively. ChAT expression in the DG was significantly lower in the NRSG than in the CG and CDG (p< 0.001), and the NRSDG showed a significant increase of ChAT expression in the DG compared to the NRSG (p< 0.001) (**Fig 7B**).

**Figure 7.**
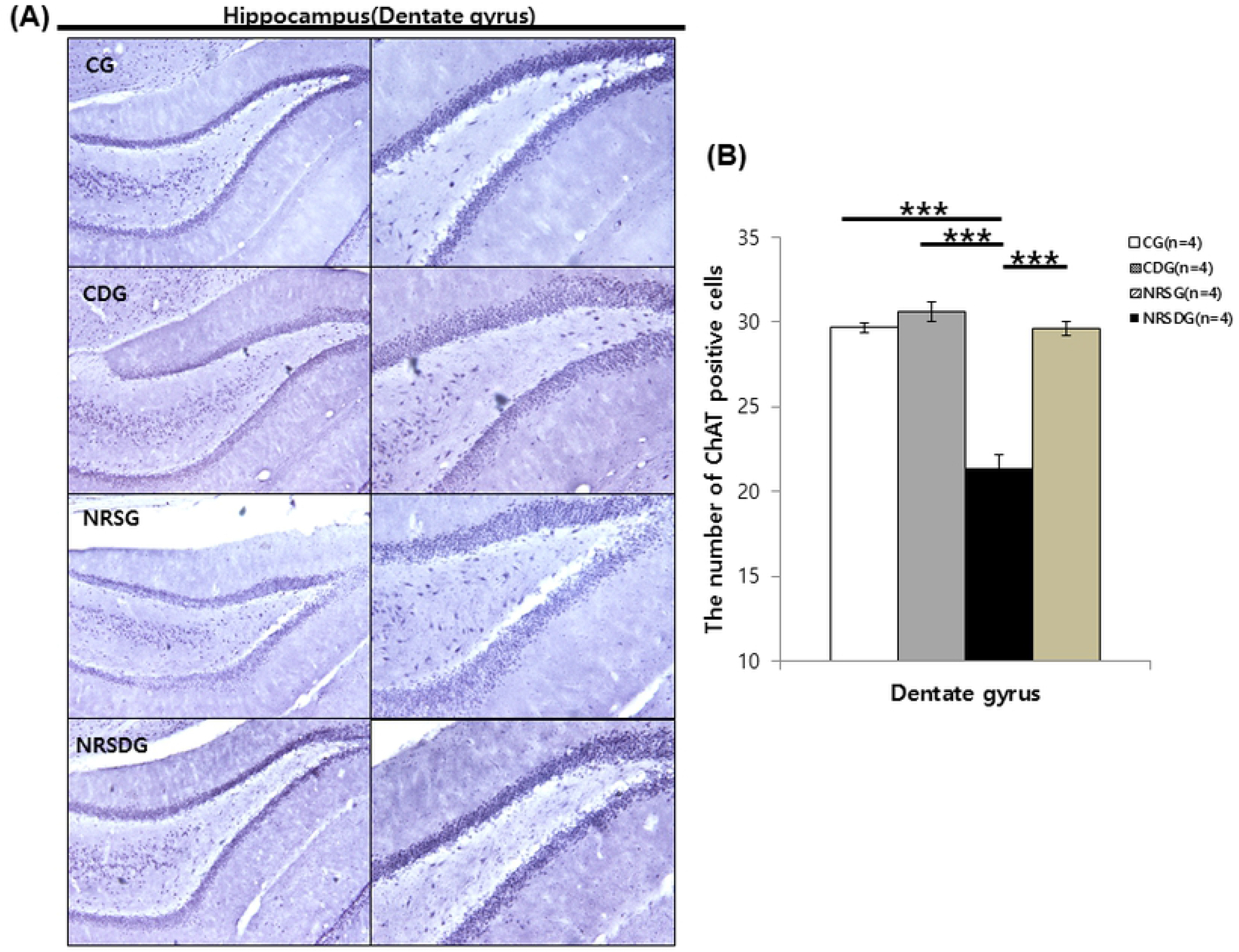
Immunohistochemisty study of the hippocampal tissue with antibodies specific for ChAT. A representative image of ChAT expression in the hippocampus in the dentate gyrus for each group (magnification X100 and X200) (A). Alpha-GPC significantly increased ChAT expression in the hippocampus of the NRSDG compared to the NRSG (B). (Error bars indicate SEM, ANOVA, Tukey’s HSD post-hoc, *** p< 0.001) (N=4).

### 3.5. Effect of α-GPC on dual stressed rat hippocampus

Immunofluorescence was performed to evaluate the effect of α-GPC on BDNF expression in the hippocampus, especially in the dentate gyrus (DG) area (**Fig 8A**). The intensity levels of BDNF expression in CG, CDG, NRSG, and NRSDG were 5222.93 ± 752, 3966.45 ± 783, 2090.859 ± 624, and 4941.28 ± 463, respectively. BDNF expression in the DG was significantly lower in the NRSG than in the CG (p< 0.05), and the NRSDG showed a significant increase of BDNF in the DG compared to the NRSG (p< 0.05) (**Fig 8B**).

**Figure 8.**
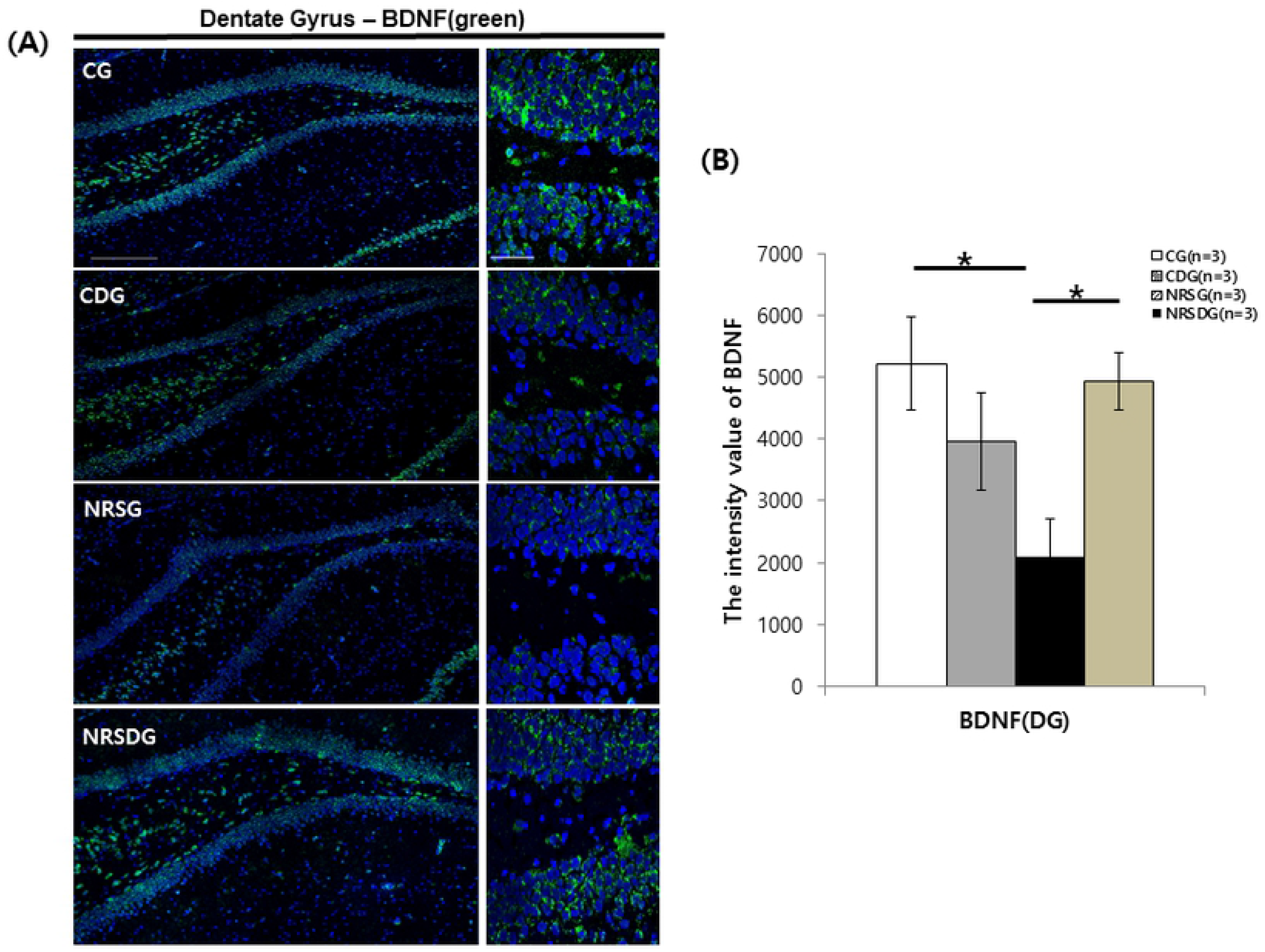
Immunofluorescence study of the hippocampal tissue with antibodies specific for BDNF. A representative image of BDNF expression in the hippocampus in the dentate gyrus for each group (magnification X200 and X400) (Scale bar= 200 and 50 μm) (A). Alpha-GPC significantly increased BDNF expression in the hippocampus of the NRSDG compared to the NRSG (B). (Error bars indicate SEM, ANOVA, LSD post-hoc, * p< 0.05) (N=3).

### 3.6. Neurogenesis assessment in neuroblasts

Immunofluorescence measurements in neuroblasts were performed to confirm the increase of immature neurons, using the DCX marker in DG at SGZ (**Fig 9A**). The number of DCX-positive cells in the CG, CDG, NRSG, and NRSDG were 126.04 ± 10.7, 107.30 ± 9.5, 85.65 ± 6.3, and 122.73 ± 8.9, respectively, which indicates that neurogenesis in the SGZ significantly decreased in the NRSG compared to the CG and the NRSDG (p< 0.05 for NRSG vs CG and NRSDG). Interestingly, the NRSDG showed a significant increase in DCX-positive cells compared to the CG (p< 0.05) (**Fig 9B)**.

**Figure 9.**
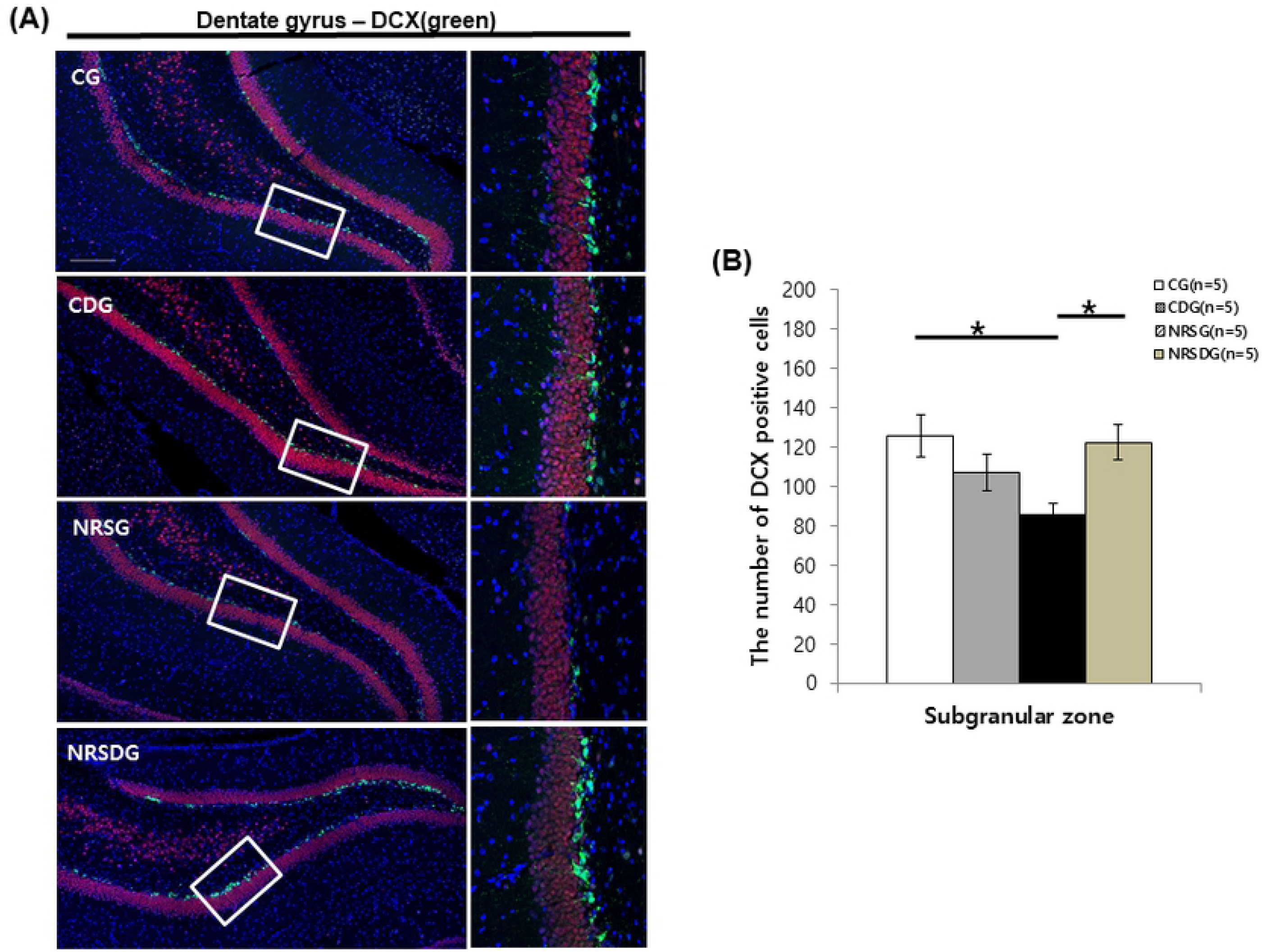
Immunofluorescence study of neuroblasts in hippocampal tissue using the DCX marker. Representative images of DCX-positive cells in the subgranular zone of the dentate gyrus in each group (magnification X200 and X400) (Scale bar= 200 and 50 μm) (A). Alpha-GPC significantly increased the number of DCX-positive cells in the NRSDG after dual stress as much as those in the CG (B). (Error bars indicate SEM, ANOVA, Tukey’s HSD post-hoc, * p< 0.05) (N=5).

## 4. Discussion

Here, we report that dual stress induced severe deficits in the behavioral performance and cholinergic activity along with neuronal degeneration in the hippocampus. While, treatment of α-GPC significantly recovered non-spatial memory impairments and it had protect to damage expression of neurotransmitters in the brain caused by dual stress in male rats. However, studies have shown that female rats are more resistant to the stress stimuli than males, demonstrating decreased behavioral performance that males showed cognitive dysfunction (19). Moreover, the estrogen that hormone affected neurogenesis to female (20). Based on these findings, we performed experiment using male rats that would be more suitable for induced model.

### 4.1. Dual stress of noise and restraint induced hearing loss, changes in organ of Corti, and elevated plasma corticosterone levels

Noise stress is significantly associated with tinnitus and cognitive dysfunction on Morris water maze test, which is the result of direct neuronal injury caused by the acoustic overpressure (6, 21). Also studies have shown an increased hearing threshold after noise exposure in rats (22). In this study, our result showed increased hearing thresholds with all frequencies as shown by ABR and DPOAE levels. Our histologic study also demonstrated significant damage and degeneration of the organ of Corti. No difference was observed in hearing and the histologic changes of the organ of Corti among the two stress-induced groups of our study, which suggests that α-GPC cannot reverse hearing or cochlear damage after the dual stresses of noise and restraint.

Organisms are continually subjected to several stressors that affect numerous physiological responses by activating the hypothalamo-pituitary-adrenal (HPA) axis and related molecular signaling including release of glucocorticoids (23). Plasma corticosterone, a well-known stress marker, since other studies have demonstrated enhancement of plasma corticosterone levels after various stress stimuli (24). Our study result also showed higher plasma corticosterone levels in the NRSG and the NRSDG compared to the control groups, which indicates that our dual stress model was properly set up, although α-GPC did not appear to regulate plasma corticosterone and appeared to be involved in cognitive function without being affected by corticosterone.

### 4.2. Neuronal cells in the hippocampus were also protected by α-GPC

Stressor including restraint stress is often used to confirm molecular and behavioral effects in experimental studies and has been helped to understand the stress-related brain pathology and the changes in cognition and severe neuronal cell loss (23, 25, 26). Other experimental studies showed that neuronal histological changes and damages in the hippocampus as well as behavioral alterations after noise or restraint stress (2). In this study, we also observed that the number of hippocampal neuronal cells significantly decreased in the NRSG compared to CG, while total number of neuronal cells significantly increased in NRSDG compared to the NRSG. Interlukin-1β (IL-1β), pro-inflammatory cytokines, is known to contribute to the actions of stress (27). Other studies reported that stress stimuli increases IL-1β in the hippocampus and central administration of IL-1β produces activation of the hypothalamic-pituitary-adrenal (HPA) axis, down-regulation of hippocampal BDNF level (27). In this study, we also observed that IL-1β expression has significantly increased in the hippocampus of NRSG while it has significantly decreased in NRSDG compared to other groups.

These results support our hypothesis that the recovery of memory function after stress with α-GPC administration could be caused by inhibition of neuronal degeneration and immune response in the hippocampus. However, the details of the inflammatory response and cell loss including apoptosis should be studied in the further research. It seems to be related to their better behavioral performance in cognitive function.

### 4.3. Alpha-*GPC protected dual stress-induced memory dysfunction*

The NOR test is a well-known assessment of non-spatial memory function in rats (28). Several studies have been shown that hippocampus is essential brain for NOR test that involved in memory processing and retrieval of object memory by interacting with perirhinal cortex (29, 30). Moreover, in the rats with hippocampal lesions, impaired novelty performance in NOR test with longer interval has been reported (31). Experimental study also showed the increased firing rates and glutamate efflux in the hippocampal pyramidal neurons which means activation of hippocampal function during NOR test (30). However, other study demonstrated that rodents with stress stimuli decreased the non-spatial memory in NOR test (2).

Alpha-GPC is a semi-synthetic derivative of phosphatidylcholine that increases acetylcholine release in the hippocampus, enhancing learning and memory function and reducing structural changes following aging (32). It promotes active form of choline that is able to reach cholinergic nerve terminals which increases the synthesis, levels and release of Ach (33). Other studies have reported that α-GPC improves the Ach level in rat hippocampus, learning and memory and brain transduction mechanisms (34, 35). In rodents, the oral lethal 50 of α-GPC is greater than 10,000 mg/kg. Oral toxicity studies in rats (up to 1,000 mg/kg/day) showed symptomology consisting of several reduced responses and there are no accompanied by histopathological correlates (36). Studies show that received various α-GPC dosage up to 300mg/kg for rodents (34).

In this study, we hypothesized that α-GPC maintains acetylcholine release in the hippocampus in stressed rats and may protect cognitive function after the severe stress. In order to demonstrate our hypothesis, we performed NOR test to investigate the changes in cognitive function after stress in rats and to evaluate the effect of α-GPC in this study. Our study results showed that the DI in the NRSG significantly decreased after the stress, which indicates that cognitive dysfunction occurred after dual stress. While, the NRSDG showed increases in DI and time spent searching for new objects, which indicates increased memory function in dual stressed rats after α-GPC administration.

### 4.4. Alpha-*GPC increased Ach levels in the hippocampus*

Decrease of cholinergic neurotransmitter from basal forebrain to hippocampus is thought to be one of the factors involved in determining memory impairment in AD disease and normal aging (37). ChAT is a most suitable marker that evaluate the activity of cholinergic neurons, which is biosynthetic enzyme of acetylcholine and it has been known to regulate Ach synthesis that affected Ach levels (38). Other study showed that decreased the ChAT activity in hippocampus after repeated restraint stress and impaired the cholinergic function in patients with memory-impaired dementia (32, 39). In this study, we demonstrated that the increase of ChAT expression in NRSDG, suggesting effects of α-GPC on cholinergic neurotransmission by increasing hippocampal ChAT activity and it seems to affect Ach level. Moreover, the neurogenesis caused by cholinesterase inhibitor and Ach is possibly due to activation of muscarinic and nicotinic receptor, and inhibition of inflammation, thereby increasing BDNF production (16).

Based on our behavioral and neurotransmitter results, we moved on to evaluate BDNF expression and neurogenesis in the hippocampus of the rats after dual stress.

### 4.5. Alpha-GPC increased BDNF expression, which may be essential for memory enhancement

BDNF is a well-known neurotrophic factor that binds to the TrkB receptor at the surface of neuronal cells, and it has been shown to interact with the signaling pathway (40). In experimental studies, knockout rodents for any neurotrophins or their receptors developed severe neuronal diseases. Moreover, the BDNF signaling pathway has been shown to play a critical role in neuronal differentiation, survival, plasticity, and cognition (12, 41–43). In this study, our results showed that the expression of BDNF in the hippocampus of the NRSDG significantly increased compared to that of the NRSG, which suggests that α-GPC significantly increases BDNF expression in the hippocampus after severe stress in rats, suggesting its contribution to neurogenesis in the hippocampus after severe stress exposure.

### 4.6. Alpha-GPC promoted neurogenesis in dual stressed rats

In the adult brain, neurogenesis occurs in the SGZ of the hippocampus at DG, and this neuronal development has been known to play an important role in the learning and memory functions of the hippocampus (6). It has been reported that immature neurons could be used to detect or process novel stimuli and new neurons at adulthood may be related to temporary storage of information in hippocampal function (20). However, studies have shown that rodents exposed to various types of stress inhibited development of neuronal progenitor cells, resulting in decreased neuroblast production in the hippocampus (28, 44, 45). Our study results also showed that the number of DCX-positive cells, a marker for neuroblasts, was significantly increased in NRSDG compared to the NRSG. Our study also supported α-GPC increased new neuronal memory function in NRSDG and these results support our hypothesis that α-GPC activates or promotes neurogenesis in the stressed hippocampus.

## 5. Conclusions

In this study, we showed that α-GPC increased ChAT and BDNF expression in the hippocampus of rats after dual stress, resulting in increased neurogenesis and cognitive function. Therefore, the mechanism by which α-GPC improves cognitive function after severe stress might involve neurogenesis and inhibit neuronal inflammation in the hippocampus. Our basic study results support a clinical role of α-GPC for patients with memory impairment after severe stress, although further studies will be needed to test its role in this area.

## Abbreviations

ABR: auditory brainstem response;
Ach: acetylcholine;
BDNF: brain-derived neurotrophic factor;
ChAT: Choline acetyltransferase;
DCX: doublecortin;
DG: dentate gyrus;
DI: discrimination index;
GM: geometric mean;
LM: light microscopy;
NOR: novel object recognition;
OC: organ of Corti;
DPOAE: distortion product otoacoustic emission;
SGZ: subgranular zone;
SPL: sound pressure level;
α-GPC: choline alphoscerate;

## Acknowledgements

This study was supported by research grants from the Basic Science Research Program of the National Research Foundation of Korea funded by the Ministry of Education, Science, and Technology (NRF2018R1D1A1A02048972) & DAEWOONG BIO (South Korea) (DWBIO/CMC_2016-0241).

